# Widespread AAV-mediated CSPG digestion impairs functional recovery after cervical spinal cord injury

**DOI:** 10.64898/2026.05.22.727165

**Authors:** Mariajose Metcalfe, Oswald Steward, Daisy Gallardo

## Abstract

Spinal cord injury (SCI) disrupts long-distance communication between the brain and spinal circuits, resulting in persistent motor, sensory and autonomic dysfunction^1^. These deficits arise from the limited regenerative capacity of adult central nervous system (CNS) neurons and the presence of a growth-inhibitory extracellular environment. Among pathways that control voluntary movement, failure of corticospinal tract (CST) regeneration is a major contributor to impaired motor function.

Strategies to promote regeneration have focused on enhancing the intrinsic growth capacity of injured neurons, such as through activation of the mTOR pathway via phosphatase and tensin homolog (PTEN) suppression, as well as reducing extrinsic inhibition through enzymatic digestion of chondroitin sulfate proteoglycans (CSPGs) using chondroitinase ABC (chABC). Because these mechanisms act through distinct but complementary processes, we investigated their combination as a strategy to improve regeneration.

Adeno-associated viral (AAV) vectors are widely used to enhance intrinsic growth pathways and represent a clinically relevant platform for gene delivery. In contrast, CSPG digestion has primarily been achieved using lentiviral or focal delivery approaches. We therefore examined whether reducing extrinsic inhibition could be implemented using AAV2-mediated chABC delivery, alone and in combination with AAV2-retro-mediated PTEN knockdown, following cervical SCI.

AAV2-mediated delivery of chABC produced robust and persistent CSPG digestion that extended beyond the injection site, and this spatial extent was influenced by viral dose and expression magnitude. Despite effective CSPG degradation, AAV2-chABC treatment did not improve functional outcomes relative to controls and did not enhance the effects of intrinsic growth activation via PTEN knockdown. Instead, AAV2-chABC treatment, alone or in combination with AAV2-retro-mediated PTEN knockdown, was associated with impaired motor performance in behavioral assays.

These findings indicate that the extent and persistence of CSPG degradation critically shape functional outcomes after SCI and that simultaneous enhancement of intrinsic growth capacity and extracellular permissiveness does not necessarily translate into improved functional recovery. Together, these results underscore the importance of carefully controlling transgene expression levels and duration in AAV-based gene therapies, where suboptimal delivery parameters may offset the benefits of otherwise promising targets.

## INTRODUCTION

Spinal cord injury (SCI) disrupts long-distance communication between the brain and spinal circuits, resulting in persistent motor, sensory and autonomic dysfunction^1^. Damage to descending motor pathways, particularly the corticospinal tract (CST), is a major contributor to impaired voluntary movement^2^, making CST regeneration a central goal of regenerative strategies following SCI^3^. Despite substantial advances in our understanding of axonal growth mechanisms, re-establishing functional connectivity across the lesion and over long distances in the adult CNS remains a major unresolved challenge.

Two broad classes of mechanisms limit axonal regeneration after SCI. These include a reduced intrinsic growth capacity within adult CNS neurons and a persistently inhibitory extracellular environment at the site of injury^4^. These barriers operate through distinct but complementary processes, in which intrinsic pathways regulate the ability of neurons to initiate and sustain growth^5–7^, whereas extrinsic factors impose inhibitory constraints within the lesion environment^8–10^. Both have therefore been targeted independently to promote regeneration.

Among these, reduced intrinsic growth capacity has emerged as a target for regenerative interventions, in part because adult CNS neurons fail to mount an effective growth response after injury. Following target innervation and circuit assembly, CNS neurons adopt a mature state optimized for synaptic stability rather than long-distance axon extension^11,12^, a transition associated with repression of growth-associated transcriptional programs^13^ and reduced activation of regeneration-associated genes (RAG)^14,15^. Together, these changes underlie the limited regenerative response of adult CNS neurons.

Restoring intrinsic growth capacity therefore relies on reactivating components of these latent developmental programs. Several RAGs promote axonal outgrowth when overexpressed, including Krüppel-like factor 7 (KLF7)^16^, signal transducer and activator of transcription 3 (STAT3)^17^ and neuronal calcium sensor-1 (NCS-1)^18^. These factors act in part by re-engaging growth-associated transcriptional networks. Among these approaches, activation of the mTOR pathway through deletion or knockdown of phosphatase and tensin homolog (PTEN)^19–23^ has produced some of the most robust CST regeneration reported to date^24,25^. PTEN suppression represents a particularly potent strategy, as it relieves a key intrinsic brake on growth signaling and enables sustained activation of pro-growth pathways^19,21,26,27^. Nevertheless, even with strong activation of intrinsic growth programs, regenerating axons frequently fail to extend beyond the immediate lesion site or achieve long-distance projections, suggesting that reactivation of intrinsic growth alone may be insufficient to fully overcome the inhibitory cues present in the injured spinal cord.

Following SCI, tissue damage triggers a cascade of cellular and molecular responses that actively reshape the lesion environment into a non-permissive state for axonal growth^28^. Cell death, inflammation and the activation of astrocytes and other glial populations drive the formation of a dense glial scar and extensive remodeling of the extracellular matrix^29–31^. These coordinated responses generate a range of inhibitory molecules derived from both damaged tissue and reactive cell populations. The injured spinal cord contains multiple sources of extrinsic inhibition, including myelin-associated inhibitors derived from damaged oligodendrocytes and myelin debris^32–35^, repulsive guidance cues expressed by reactive cells^36^, and extracellular matrix components enriched within the glial scar^37^. Among these, chondroitin sulfate proteoglycans (CSPGs), produced primarily by reactive astrocytes and other glial cells, constitute a major component of the glial scar^38^ and accumulate rapidly after injury^39^. CSPGs restrict axonal extension through both biochemical and physical mechanisms, including receptor-mediated inhibition of growth signaling and the formation of the glial scar that acts as a physical barrier to axon extension^40^.

Accordingly, targeting CSPGs has emerged as a central strategy to modulate the inhibitory extracellular environment. Enzymatic digestion of CSPGs using chondroitinase ABC (chABC) enhances axonal sprouting and plasticity across multiple injury models^41–44^, supporting the concept that removal of extracellular inhibition can facilitate regeneration. However, modulation of the extracellular environment alone does not fully restore long-distance axonal growth, as regenerating axons often fail to sustain elongation in the absence of intrinsic growth activation programs. For this reason, combinatorial strategies targeting both intrinsic and extrinsic factors have been widely proposed as a rational approach to promote regeneration after SCI^45–49^. Conceptually, simultaneously enhancing neuronal growth capacity while reducing environmental inhibition should create a permissive state for axonal extension.

Given the reproducible effects of CSPG degradation on axonal sprouting and plasticity, a logical next step is to ask whether this strategy can be translated into a clinically viable gene delivery platform, particularly in the context of combinatorial approaches that already target intrinsic growth pathways. Prior studies of CSPG degradation have relied on focal delivery of bacterial chABC^50,51^, which produces transient enzymatic activity in vivo and therefore would require repeated administrations to maintain CSPG digestion, or lentiviral expression systems^52^, which, while enabling sustained expression, face significant translational limitations due to their genomic integration, insertional mutagenesis risk and regulatory barriers that have limited their clinical adoption relative to non-integrating vectors^53–55^. In parallel, strategies designed to enhance intrinsic neuronal growth capacity, such as targeting PTEN/mTOR signaling, are most commonly delivered using adeno-associated viral (AAV) vectors^19,21,22,24,25,56–58^, establishing AAV as the prevailing platform for modulating neuron-intrinsic programs in vivo. These considerations raise a practical and translational question: can CSPG degradation be deployed using the same AAV-based framework to enable unified targeting of both intrinsic and extrinsic barriers?

AAV vectors offer the potential for sustained, widespread and clinically translatable transgene expression^59,60^, making them an attractive platform for implementing extracellular matrix modulation alongside intrinsic growth activation. Critically, the development of retrogradely transported AAV variants (AAV-retro) enables efficient genetic manipulation of supraspinal neurons through spinal cord delivery alone^61,62^, achieving greater than 90% transduction efficiency in corticospinal and other supraspinal populations without requiring direct intracranial delivery. This is particularly relevant in the context of SCI, where spinal injections can simultaneously target multiple supraspinal populations whose projections are interrupted by injury, across a clinically accessible route^56,63^. Of particular translational relevance, combining AAV2-mediated spinal chABC delivery with AAV-retro-mediated corticospinal PTEN knockdown permits coordinated modulation of the extracellular environment and neuronal intrinsic growth programs through a single cocktail of AAV vectors administered in any proportion desired, a combination strategy that could be advanced as a unified therapeutic agent in clinical translation.

In this study, we examined whether AAV2-mediated delivery of chABC could serve as a platform for sustained CSPG degradation and whether this approach enhances functional recovery after cervical SCI, alone and in combination with AAV2-retro-mediated PTEN knockdown. We further investigated how the extent and spatial distribution of AAV-driven transgene expression shape functional outcomes, with direct implications for the design of combined gene therapy strategies targeting both intrinsic and extrinsic barriers to repair.

## RESULTS

### AAV-mediated chABC delivery produces widespread and sustained CSPG digestion

To determine whether AAV can deliver functionally active chABC in the adult spinal cord, we first evaluated the feasibility of AAV-mediated chABC expression. Because the chABC coding sequence approaches the packaging limits of AAV vectors^64^, we utilized a previously described codon-optimized chABC sequence^51^, incorporated into a compact expression cassette under the PGK promoter (Supplementary Fig. 1).

To assess functional enzymatic activity in vivo, adult rats were injected at cervical level 5 (C5) with AAVrh10-chABC or AAV2-chABC, with direct injection of bacterial chABC included as a reference condition. CSPG digestion was evaluated by immunofluorescence detection of chondroitin-4-sulfate (C-4-S), an epitope exposed following enzymatic cleavage of chondroitin sulfate glycosaminoglycan chains^52,65^.

AAV-GFP-injected controls showed no detectable C-4-S immunoreactivity, confirming the absence of baseline CSPG digestion (Figure 1A). Bacterial chABC produced localized C-4-S signal restricted to the vicinity of the injection site (Figure 1B), consistent with previously reported focal enzyme delivery^50,51^. In contrast, both AAVrh10-chABC and AAV2-chABC produced robust C-4-S immunoreactivity at the injection site (Figure 1C-D), indicating effective expression and enzymatic activity following AAV-mediated gene transfer. Strikingly, CSPG digestion extended far beyond the injection site, with C-4-S immunoreactivity detected in thoracic spinal cord 1 cm caudal to the injection (Figure 1E-F) and persisting in lumbar spinal segments 2 cm caudal (Figure 1G-H).

**Figure 1.**
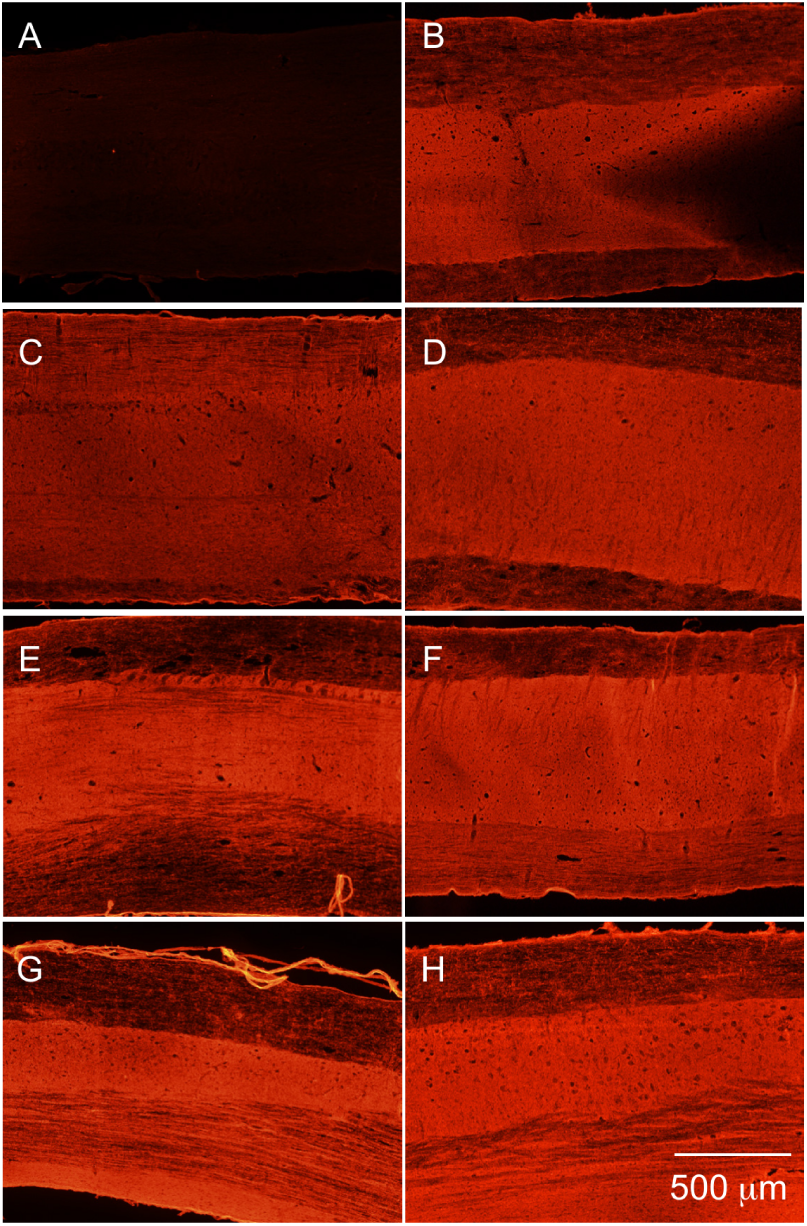
AAV-mediated chABC expression produces widespread CSPG digestion across spinal cord segments. CSPG digestion was assessed by immunofluorescence detection of chondroitin-4-sulfate (C-4-S), a marker of chondroitin sulfate proteoglycan cleavage. (A-B) Control conditions. (A) AAV2-GFP shows no detectable C-4-S immunoreactivity. (B) Bacterial chABC produces localized CSPG digestion restricted to the injection site. (C-D) Cervical spinal cord (injection site, C5). AAVrh10-chABC (C) and AAV2-chABC (D) produce robust C-4-S immunoreactivity at the site of delivery. (E-F) Thoracic spinal cord (1 cm caudal to injection). C-4-S immunoreactivity is detected following AAVrh10-chABC (E) and AAV2-chABC (F), indicating extension of enzymatic activity beyond the injection site. (G-H) Lumbar spinal cord (2 cm caudal to injection). C-4-S immunoreactivity persists in distal spinal segments following AAVrh10-chABC (G) and AAV2-chABC (H), demonstrating multi-segmental distribution of CSPG digestion. Scale bar, 500 µm.

Together, these data establish that AAV-mediated delivery produces functional and spatially extensive CSPG digestion in the spinal cord, providing a platform to modulate the inhibitory extracellular environment in vivo.

### AAV2-mediated CSPG digestion is sustained after injury and tunable by viral dose

Having established functional and spatially extensive AAV-mediated chABC activity, we next defined its temporal profile and dose dependence following SCI. Based on initial serotype comparisons, AAV2 was selected for all subsequent experiments to align with AAV-based strategies targeting intrinsic growth pathways and support a consistent combinatorial platform in which AAV2-chABC addresses extrinsic inhibition and AAV2-retro-shPTEN targets corticospinal neurons. CSPG digestion was assessed by immunodetection of C-4-S in the injured spinal cord following AAV2-chABC delivery at the time of injury.

Robust C-4-S immunoreactivity was detected at the lesion site as early as 2 days post-injection (Fig. 2A) and remained detectable at 7, 14 and 21 days post-injection (Fig. 2B-D), indicating that AAV2-mediated chABC expression produces sustained CSPG digestion in the injured spinal cord.

**Figure 2.**
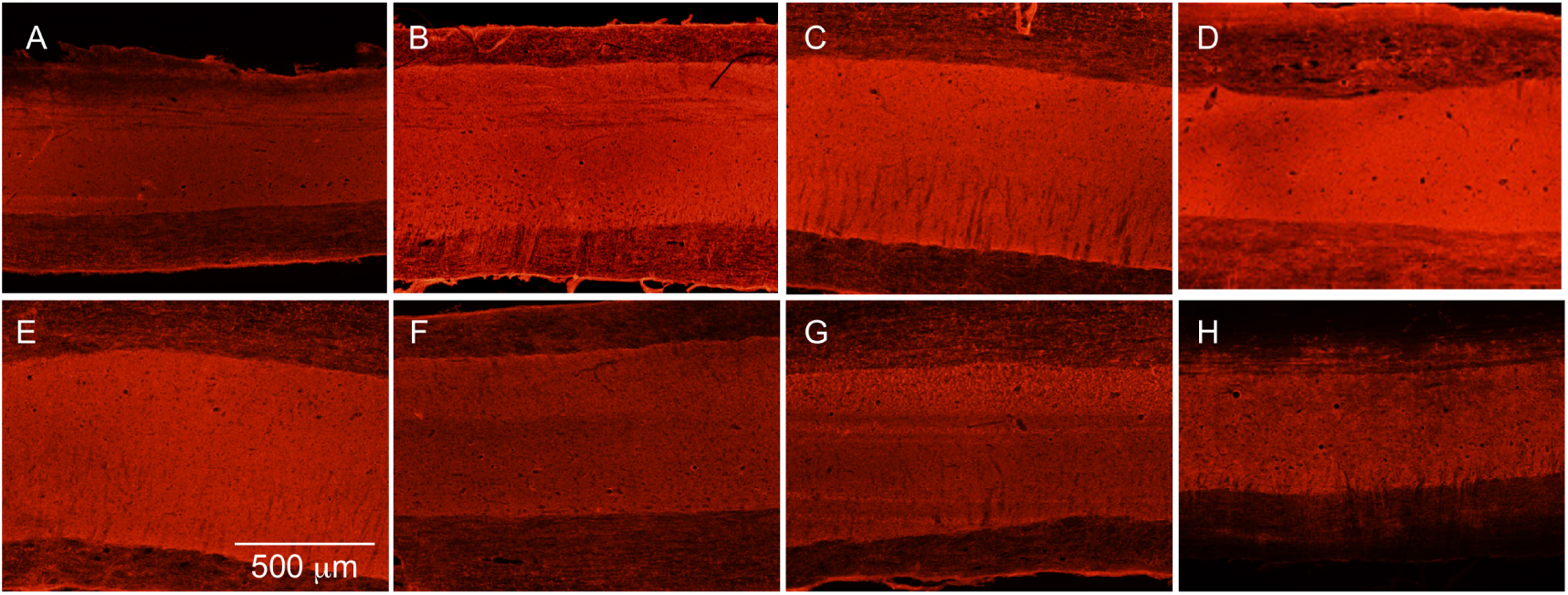
Temporal and dose-dependent dynamics of AAV2-mediated chABC activity in the spinal cord. (A-D) Representative images of C-4-S immunoreactivity at 2, 7, 14, and 21 days post-injection (dpi) following AAV2-chABC delivery at 1.8E10 GC/animal, demonstrating sustained CSPG digestion over time. (E-H) Representative images of C-4-S immunoreactivity at 14 dpi across decreasing AAV2-chABC doses: 1.8E10 (E), 9.0E9 (F), 1.8E9 (G), or 3.6E8 (H) GC/animal. Lower doses produce more spatially restricted CSPG digestion. Scale bar, 500 µm.

To determine whether the spatial extent of digestion could be modulated, animals received AAV2-chABC across a range of doses and were assessed at 14 days post-injection. Higher doses resulted in broader and more uniform C-4-S immunoreactivity extending across multiple spinal cord segments (Fig. 2E-F), whereas lower doses produced progressively more spatially restricted patterns of CSPG digestion (Fig. 2G-H), demonstrating dose-dependent control over the extent of extracellular matrix remodeling.

These findings establish that AAV2-mediated chABC activity is sustained after SCI and that its spatial extent can be titrated through viral dose, providing a tunable platform for engaging inhibitory cues across the injury environment.

### AAV2-chABC and AAV2-retro-shPTEN achieve effective target engagement in the injured spinal cord

Building on these observations, we next combined AAV2-mediated CSPG degradation with targeted activation of intrinsic growth pathways. Intrinsic growth capacity was enhanced via AAV2-retro-mediated delivery of shRNA targeting PTEN^25,57^ (AAV2-retro/shPTEN), a well-established regulator of the mTOR pathway^22^, whose suppression promotes axonal growth in CST neurons^25,56^. AAV2-retro efficiently transduces supraspinal neurons following intraspinal delivery, including CST neurons and other descending pathways^56,63^, providing access to neuronal populations that project to the spinal cord without requiring intracranial injections.

Animals received intraspinal AAV2-chABC at the time of injury, either alone or in combination with AAV2-retro/shPTEN (Combo), allowing concurrent modulation of extracellular inhibitory cues at the lesion site and intrinsic growth programs in supraspinal neurons through a shared intraspinal delivery route. Control animals received AAV2-retro-GFP. Animals were euthanized 14 days post-injury (dpi) and CSPG digestion was assessed by C-4-S immunodetection.

In AAV2-GFP control animals, no C-4-S immunoreactivity was observed in uninjured spinal cord, with signal largely restricted to the lesion site following SCI (Fig. 3A-B). AAV2-chABC delivery induced widespread C-4-S immunoreactivity in uninjured tissue (Fig. 3C) and following SCI, C-4-S signal increased and redistributed around the lesion site (Fig. 3D), indicating that the extent and spatial pattern of CSPG degradation was influenced by injury context.

**Figure 3.**
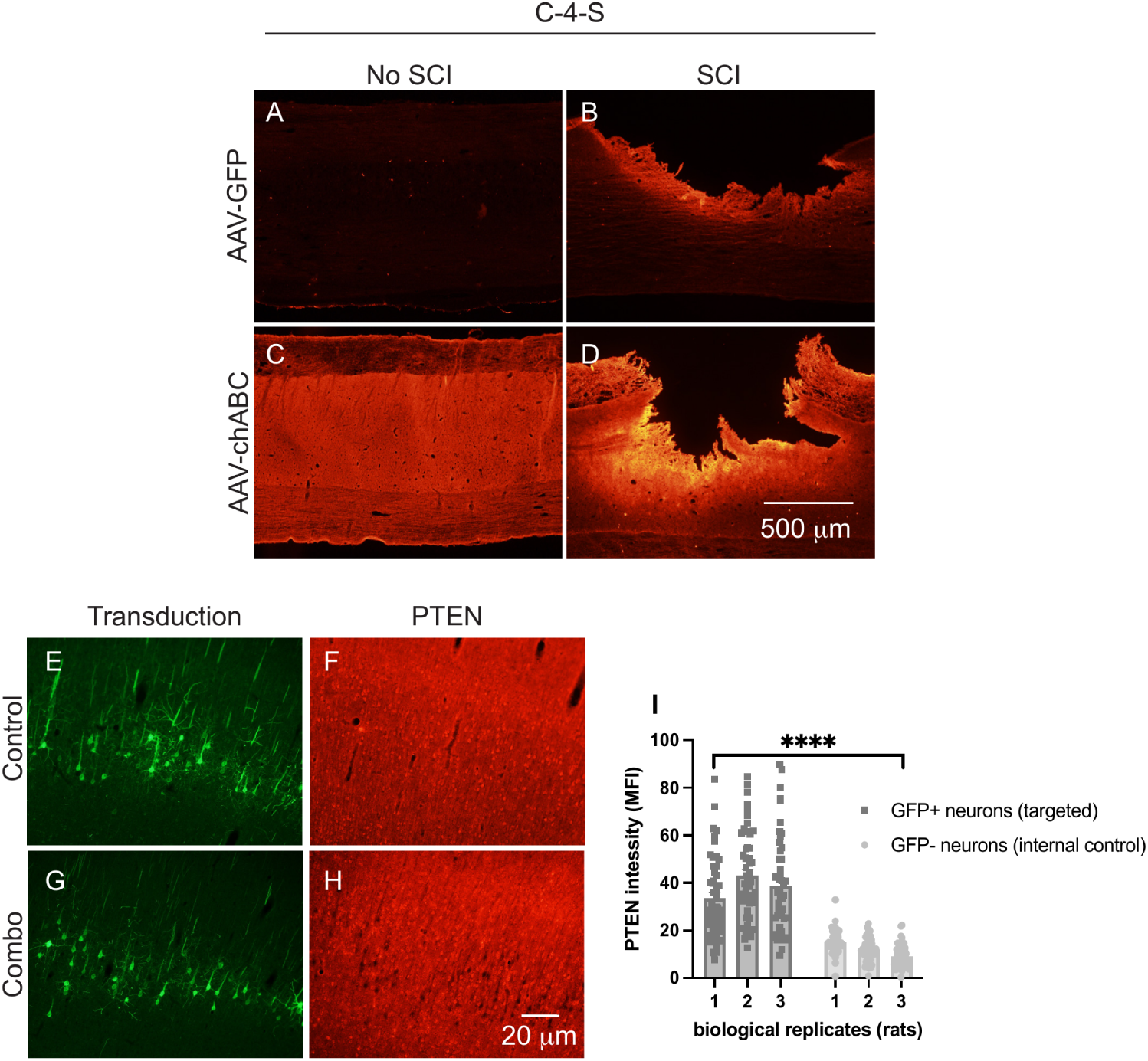
AAV2-mediated spinal CSPG degradation and cortical PTEN knockdown confirm target engagement. Representative images of C-4-S immunoreactivity in spinal cord sections following AAV delivery in the absence (No SCI) or presence of cervical spinal cord injury (SCI). (A-B) AAV2-GFP controls show minimal C-4-S signal in uninjured tissue (A) and increased signal localized to the lesion site following SCI (B). (C-D) AAV2-chABC induces widespread CSPG degradation in uninjured spinal cord (C), which is further enhanced and redistributed in the presence of injury (D), indicating that the spatial pattern of CSPG digestion is influenced by injury context. (E-H) Validation of PTEN knockdown in motor cortex neurons. GFP fluorescence confirms efficient neuronal transduction under control (E) and combination treatment conditions (G). PTEN immunostaining shows preserved expression in GFP control animals (F) and reduced signal in GFP+ neurons following AAV2-retro-shPTEN delivery (H). (I) Quantification of PTEN immunofluorescence intensity (MFI) in GFP+ (transduced) and neighboring GFP- (non-transduced) neurons across biological replicates (n = 3 animals; 50 neurons per condition per animal). GFP+ neurons exhibit significantly reduced PTEN signal compared to GFP-neurons (linear mixed-effects model; p < 0.0001). Data are presented as mean ± SEM. Scale bars, 500 µm (A-D), 20 µm (E-H).

Engagement of intrinsic growth pathways was assessed by evaluating PTEN knockdown in corticospinal neurons^25,63^ following intraspinal delivery of AAV2-retro/shPTEN. GFP fluorescence confirmed efficient transduction of motor cortex neurons under both control and combination treatment conditions (Fig. 3E, G). PTEN immunostaining revealed preserved expression in GFP control animals, whereas GFP+ neurons in shPTEN-treated animals exhibited markedly reduced PTEN signal (Fig. 3F, H). Quantification of PTEN immunofluorescence intensity confirmed a significant reduction in GFP+ transduced neurons compared to neighboring GFP- neurons across biological replicates (Fig. 3I; p < 0.0001), with adjacent GFP- layer V neurons serving as within-section internal controls.

These results confirm effective target engagement of both extracellular inhibitory cues and intrinsic growth pathways at the time of functional assessment.

### AAV2-chABC treatment impairs functional recovery after cervical SCI independently of PTEN knockdown

Having established that both intrinsic and extrinsic pathways can be robustly engaged by effective PTEN knockdown and CSPG degradation respectively, we next assessed the functional consequences of these interventions following cervical SCI. Because C5 dorsal hemisection disrupts CST projections critical for skilled forelimb movement^66^, functional recovery was evaluated using the staircase reaching task^67,68^, which is sensitive to CST-dependent motor control. All groups exhibited a marked decline in performance immediately after injury, followed by partial recovery over time (Fig. 4C).

**Figure 4.**
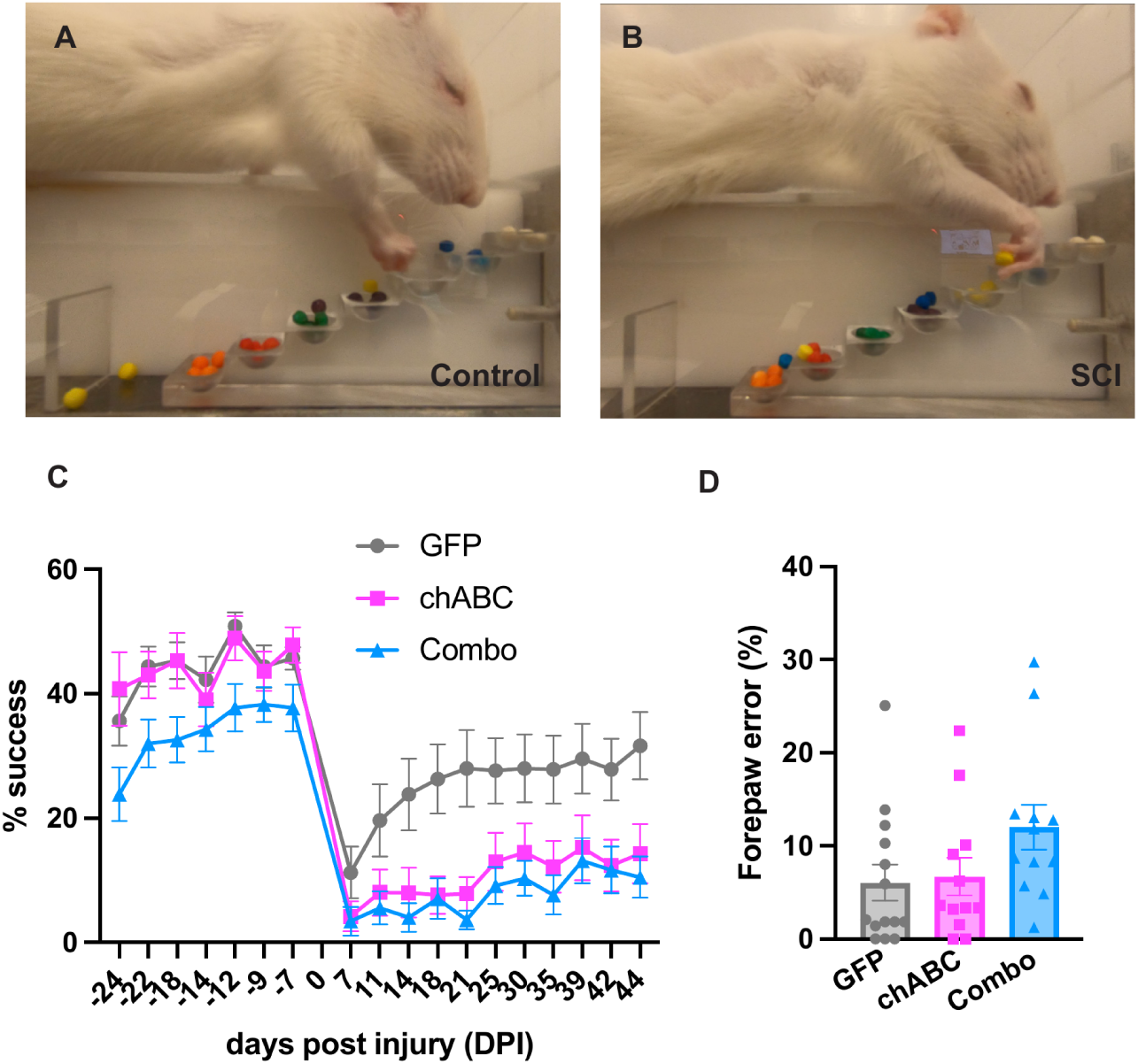
AAV2-chABC treatment, alone and in combination with PTEN knockdown, impairs functional recovery after cervical spinal cord injury. (A-B) Representative images from the staircase pellet reaching task in uninjured control animals (A) and following cervical SCI (B). (C) Quantification of pellet retrieval success over time. Animals received AAV2-retro-GFP (GFP), AAV2-chABC (chABC), or combined treatment (Combo: AAV2-chABC + AAV2-retro-shPTEN/GFP). Both chABC- and combination-treated animals exhibit reduced recovery compared to GFP controls, with a more pronounced and consistent effect in the combination group (two-way repeated measures ANOVA, significant main effect of treatment, p = 0.0010; no significant interaction between time and treatment). Time-resolved post hoc comparisons are provided in Supplementary Fig. 2. (D) Forepaw placement errors in the ladder rung task at 30 dpi. Error rates were not significantly different across groups (one-way ANOVA, Tukey’s multiple comparisons test, p = 0.1102), although the combination group shows a modest increase in errors consistent with panel C. Data are presented as mean ± SEM (n = 15 GFP, n = 15 chABC, n = 15 Combo).

AAV2-chABC-treated animals, alone or in combination with AAV2-retro-shPTEN (Combo), did not show improved recovery relative to GFP controls. Rather than improving recovery, both treatment groups performed worse than GFP controls over the observation period (Fig. 4C; two-way repeated measures ANOVA, significant main effect of treatment, p = 0.0010). Post hoc comparisons revealed no differences between groups at early timepoints (7 and 11 dpi). Beginning at 14 dpi, the combination group exhibited reduced performance relative to GFP controls, while AAV2-chABC-treated animals did not differ significantly at that time point. At later time points (18-44 dpi), both AAV2-chABC and combination groups showed persistently reduced performance compared to GFP controls, with more consistent deficits observed in the combination group (Supplementary Fig. 2).

Forelimb performance was further assessed using the ladder rung task^25^ at 30 dpi, when performance had stabilized across groups. Forepaw placement errors were not significantly different across groups (Fig. 4D; p = 0.1102), although the combination group showed a modest increase in errors, consistent with the staircase reaching findings.

These results indicate that AAV2-chABC treatment, alone or combined with PTEN knockdown, was associated with impaired motor performance relative to controls, and that simultaneous modulation of extracellular inhibitory cues and intrinsic growth programs did not improve functional recovery after SCI.

## DISCUSSION

AAV2-mediated CSPG degradation, either alone or in combination with PTEN knockdown, was associated with reduced performance relative to GFP controls. This active impairment, rather than merely a failure to improve, is an important distinction as it indicates that the observed functional deficits reflect a consequence of the intervention rather than insufficient statistical power or inadequate target engagement. Indeed, both CSPG degradation and PTEN knockdown were robustly achieved, yet neither alone nor in combination produced functional benefit.

These findings contrast with prior studies using bacterial or lentiviral chABC delivery, which reported functional improvements alongside digestion largely confined to the lesion epicenter and immediate surrounding segments^39,50,65,69,70^, in contrast to the multi-segmental distribution observed here (Figs. 1, 2). A key difference between those approaches and the present study lies in the delivery platform and the resulting spatial extent of CSPG digestion. Notably, lentiviral vectors produce more spatially restricted transgene expression than AAV vectors due to differences in particle size^71^ and spread from the injection, which may partly account for the more localized digestion profiles reported with lentiviral delivery. This difference in the spatial and temporal profile of CSPG degradation may underlie the divergent functional outcomes.

CSPGs are key extracellular matrix components that contribute to structural stability and restrict plasticity in the adult CNS^72,73^, in part through their role in perineuronal nets (PNNs)^74^, which regulate synaptic organization and circuit refinement. While transient reduction of CSPG-mediated inhibition can promote plasticity, widespread or sustained disruption of these structures may compromise synaptic stability and functional connectivity. Although long-term chABC expression has not shown negative effects^65^, and enzymatic removal of PNNs using chABC after SCI can reopen a window of plasticity that improves motor function independent of glial scar breakdown^43^, permanent degradation of PNNs worsens motor outcomes in rats^75^. Taken together, these observations suggest that CSPG digestion confined to the lesion environment supports functional recovery, whereas pan-digestion extending into the uninjured spinal cord, where CSPGs contribute to normal circuit function, may be detrimental. Given the widespread and sustained CSPG degradation observed here, it is possible that prolonged extracellular matrix modification disrupted the balance between plasticity and circuit stabilization required for effective functional recovery^76,77^.

The dose-response data presented here indicate that the spatial extent of CSPG digestion can be modulated by adjusting viral titer, with lower titers producing a more spatially restricted CSPG degradation. The high titer used in the functional experiments was selected to maximize target engagement, based on the premise that more complete removal of inhibitory cues would be beneficial. The impaired outcomes observed suggest that this premise warrants re-examination. Consistent with findings showing that regulated and temporally controlled lentiviral-driven chABC expression is required for optimal functional recovery^69^, these results highlight that constraining both the spatial extent and duration of CSPG digestion to the lesion environment may be critical parameters for future studies.

More broadly, these findings suggest that simultaneous and widespread removal of inhibitory constraints may be detrimental to functional recovery, and that successful repair may require spatial and temporal regulation of growth-promoting interventions. A staged approach, in which initial growth and plasticity are followed by refinement and stabilization of functional connections, may be more effective than continuous or widespread transgene expression. Consistent with this, studies targeting Nogo-A, a myelin-associated inhibitor of axonal growth in the adult CNS^78^, have demonstrated the importance of sequential interventions, in which axonal growth precedes the selection and pruning of meaningful connections^79,80^. These observations support a model in which recovery requires not only promoting axonal growth but also preserving appropriate inhibitory control to enable effective circuit integration.

AAV vectors carry well-established advantages for clinical translation, including manufacturing scalability and regulatory precedent^81^. However, our findings indicate that safety remains a critical challenge, as widespread and sustained modulation of growth-promoting pathways may lead to unintended functional consequences. From a translational perspective, these results therefore highlight critical design considerations for AAV-based gene therapy, including the need for precise control over the spatial distribution and duration of transgene expression, addressing issues of targeting specificity and safety^82–84^.

Future studies will determine whether temporally controlled expression systems, such as inducible AAV platforms^69^, can better balance plasticity and circuit stability by allowing expression to be titrated or discontinued once therapeutic goals are achieved. Delayed delivery paradigms may provide an opportunity to harness plasticity at stages when circuit reorganization is more favorable, as demonstrated in other injury models^85^. Such approaches may help refine strategies that promote recovery while avoiding the detrimental effects of sustained transgene expression as observed here.

Collectively, these findings underscore that functional recovery after SCI depends not only on engaging the right molecular targets, but on precisely controlling the extent, timing and location of those interventions within the injured CNS.

## MATERIALS AND METHODS

### Animals

All procedures involving animals were approved by the Institutional Animal Care and Use Committee (IACUC) at the University of California, Irvine and were conducted in accordance with NIH guidelines for the care and use of laboratory animals. Adult Sprague-Dawley rats were used for all experiments. Animals were housed in a temperature- and humidity-controlled vivarium on a 12h light/dark cycle with access to food and water. Animals were randomly assigned to experimental groups, and all analyses were performed by investigators blinded to treatment conditions.

### AAV-vector design and chABC construct generation

A codon-optimized chondroitinase ABC (chABC) sequence previously described^51^ was received as a gift from Dr. Elizabeth Muir and was used for all AAV constructs. The chABC coding sequence was cloned into a compact AAV expression cassette under the control of the PGK promoter, followed by a human growth hormone (hGH) polyadenylation signal. The final construct was designed to remain within AAV packaging constraints^64^ and vector integrity was confirmed by sequencing. AAV vectors were produced by the University of Pennsylvania Viral Vector Core.

### AAV-mediated chABC delivery in rat spinal cord

Adult rats (Sprague-Dawley, female, n = 20; 8-10 weeks old at time of surgery) received intraspinal injections at cervical level 5 (C5). AAV vectors encoding chABC were injected at four sites within the cervical spinal cord positioned 0.5 mm rostral and 0.5 mm caudal to the injury site, using coordinates previously validated for lentiviral chABC delivery^39^. Control animals received AAV-GFP injections (n = 5) using the same coordinates. For comparison with focal enzyme delivery, a separate group received bacterial chABC injections (n = 5) at the same spinal level as previously described^50^. Two AAV serotypes were evaluated, AAVrh10-chABC (n = 5) and AAV2-chABC (n = 5) (PGK promoter, 1.8E10 GC/animal, 0.3 µl per injection). After the designated post-injection interval (3 or 7 weeks post-injection) animals were transcardially perfused with PBS followed by 4% paraformaldehyde (PFA). Spinal cords were collected, post-fixed, cryoprotected, and sectioned longitudinally at 30 µm.

### Detection of CSPG digestion

CSPG digestion was assessed by immunofluorescence for chondroitin-4-sulfate (C-4-S), an epitope exposed following chondroitinase-mediated cleavage of CS-GAG side chains. Sections were processed using standard immunofluorescence methods with anti-chondroitin-4-sulfate antibody (clone C4, MP Biomedical, Cat #0869100-CF, 1:5000; discontinued) and imaged under identical acquisition settings across all experimental groups.

### Assessment of spatial distribution of chABC activity

To evaluate the spatial extent of AAV-mediated chABC activity, C-4-S immunoreactivity was examined at the cervical injection site (C5) and at distal spinal cord segments. A 1.6 cm cervical block encompassing the injection region was imaged to assess CSPG digestion at the site of delivery. To assess long-range distribution, additional sections were collected from regions located 1 cm caudal to the injection site, corresponding to thoracic spinal cord, and 2 cm caudal to the injection site, corresponding to lumbar spinal cord. For timing studies, female rats (8-10 weeks old; n = 12 total) received AAV2-chABC injections at C5 and were perfused at 2 days (n = 3), 7 days (n = 3), 14 days (n = 3) or 21 days (n = 3) post-injection. For dose-response studies, female rats (8-10 weeks old; n = 12) received AAV2-chABC at concentrations of 1.8E10, 9.0E9, 1.8E9, or 3.6E8 GC/animal and were perfused 14 days post-injection.

### Cervical spinal cord injury and experimental groups

All functional experiments were performed in adult rats subjected to cervical level 5 (C5) dorsal hemisection injury. Immediately following injury, animals received intraspinal injections of AAV2-chABC as described above. For combination experiments (Combo), animals additionally received AAV2-retro-shPTEN delivery. Control groups included animals receiving AAV2-retro-GFP alone.

### AAV2-retro-mediated PTEN knockdown in corticospinal neurons

To achieve targeted knockdown of PTEN in corticospinal tract (CST) neurons, animals received intraspinal injections of AAV2-retro encoding shPTEN or GFP control into the cervical spinal cord at the time of injury. The retrograde tropism of this vector enables transduction of neurons projecting to the injection site, including layer V motor cortex neurons. Animals received two intraspinal injections of 0.3 µl each (1E13 GC/ml stock; 6E9 GC/animal total) of either AAV2-retro-shPTEN or AAV2-retro-GFP (Addgene #37825). For cortical validation, brains were collected at 10 weeks post-injury, post-fixed in 4% PFA, cryoprotected, and sectioned coronally at 30 µm.

### Quantification of PTEN knockdown in individual neurons

To quantify PTEN expression following AAV2-retro-mediated knockdown, mean fluorescence intensity (MFI) of PTEN immunoreactivity was measured in individual neurons using ImageJ (NIH). For each biological replicate (n = 3 animals), 50 GFP-positive (transduced) neurons were identified based on GFP fluorescence and selected for analysis. For each GFP-positive neuron, a neighboring large layer V neuron lacking GFP signal (GFP-) was selected as an internal control within the same section, under identical imaging conditions. Background signal was subtracted from each measurement prior to analysis.

### Staircase pellet-reaching task

Skilled forelimb function was assessed using the single-pellet staircase reaching task as previously described^67,68^. Animals were trained to stable performance prior to injury. Adult Sprague-Dawley rats (female, n = 50; 8-10 weeks old) were subjected to C5 dorsal hemisection and assigned to one of three experimental groups: (1) C5 dHx + AAV2-retro-GFP (n = 19), (2) C5 dHx + AAV2-chABC (1.8E10 GC/animal; n = 16), or (3) C5 dHx + AAV2-chABC + AAV2-retro-shPTEN (1.8E10 and 6E9 GC/animal, respectively; n = 15). Animals were tested three times per week through 45 dpi and performance was quantified as the percentage of successful pellet retrievals relative to total attempts. Behavioral testing was performed by experimenters blinded to treatment group.

### Ladder rung walking task

Motor coordination and forelimb placement accuracy were assessed in the same cohort using the horizontal ladder rung task, as previously described^86^. Animals were tested at 30 dpi. Error rate was calculated as the percentage of incorrect placements relative to total forepaw placements. Values from left and right forelimbs were combined to generate a single error metric per animal. Scoring was performed by experimenters blinded to treatment group.

### Statistical analysis

All data are presented as mean ± SEM. For staircase reaching data, the effects of time and treatment were assessed using a two-way repeated measures ANOVA. Time-resolved post hoc comparisons were performed using a linear mixed-effects model with Dunnett’s multiple comparisons test, with time and treatment as fixed effects and subject as a random effect. For ladder rung error analysis, group differences were assessed using one-way ANOVA followed by Tukey’s multiple comparisons test. For PTEN fluorescence quantification, individual neuron MFI values were analyzed using a linear mixed-effects model with GFP status (GFP+ vs. GFP-) as a fixed effect and animal as a random effect, accounting for the nested structure of neurons within biological replicates and avoiding pseudo-replications. Statistical significance was set at p < 0.05 for all analyses. All statistical tests were performed in GraphPad Prism v11.

## Acknowledgements

Supported by NIH grant supplement from NS047718 and NIH grant NS132773 to MM. We gratefully acknowledge the mammalian chABC sequence from Dr. Elizabeth Muir. Thanks to Ardi Gunawan for skilled animal surgery.

## SUPPLEMENTARY FIGURES

**Supplementary Figure 1.**
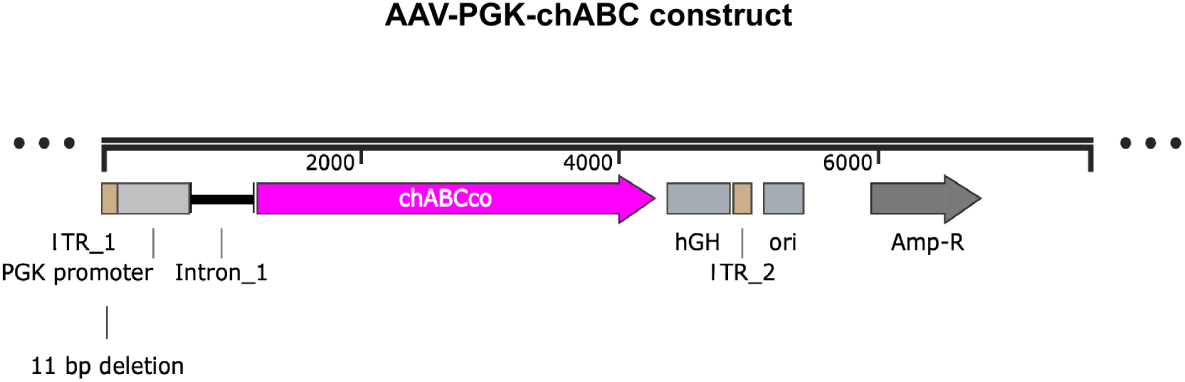
Schematic of the AAV expression cassette encoding codon-optimized chABC. Linear schematic of the AAV2 vector used in this study. The codon-optimized chABC sequence was cloned into a compact expression cassette under the control of the PGK promoter and followed by a human growth hormone (hGH) polyadenylation signal. An intron upstream of the coding sequence and flanking LTR elements are indicated, including an 11 bp deletion within the LTR. Additional plasmid backbone elements (origin of replication, ori; ampicillin resistance, AmpR) are shown for reference. Total construct length is 7,639 bp.

**Supplementary Figure 2.**
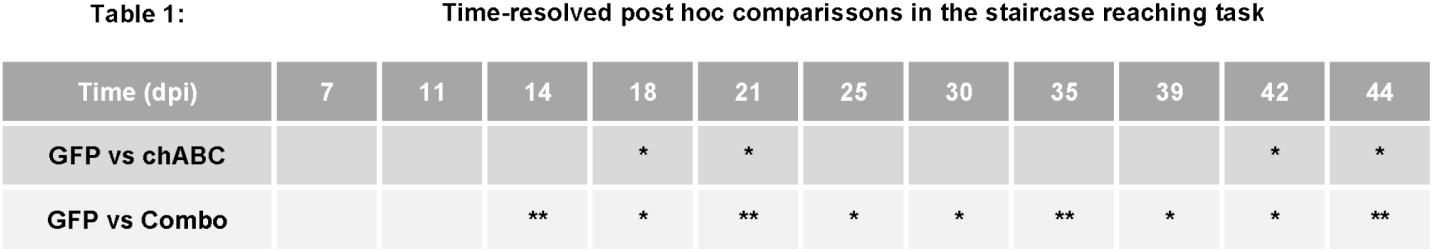
Time-resolved post hoc comparisons of staircase reaching performance following AAV-mediated interventions. Post hoc comparisons were performed using a mixed-effects model with Dunnett’s multiple comparisons test to evaluate differences between treatment groups at each time point. No significant differences were observed at early time points (7-11 dpi). Beginning at 14 dpi, the combination group exhibited reduced performance compared to GFP controls. At later time points (18-44 dpi), both AAV2-chABC and combination-treated animals showed reduced performance relative to GFP controls, with more consistent effects in the combination group. Symbols indicate significant differences relative to GFP controls (Dunnett’s test; *p < 0.05, **p < 0.01, ***p < 0.001).

## Notes

### Competing Interest Statement

The authors have declared no competing interest.

